# Single Photon smFRET. II. Application to Continuous Illumination

**DOI:** 10.1101/2022.07.20.500888

**Authors:** Ayush Saurabh, Matthew Safar, Mohamadreza Fazel, Ioannis Sgouralis, Steve Pressé

## Abstract

Here we adapt the Bayesian nonparametrics (BNP) framework presented in the first companion manuscript to analyze kinetics from single photon, single molecule Förster Resonance Energy Transfer (smFRET) traces generated under continuous illumination. Using our sampler, BNP-FRET, we learn the escape rates and the number of system states given a photon trace. We benchmark our method by analyzing a range of synthetic and experimental data. Particularly, we apply our method to simultaneously learn the number of system states and the corresponding kinetics for intrinsically disordered proteins (IDPs) using two-color FRET under varying chemical conditions. Moreover, using synthetic data, we show that our method can deduce the number of system states even when kinetics occur at timescales of interphoton intervals.

**Why It Matters:** In the first companion manuscript of this series, we developed new methods to analyze noisy smFRET data. These methods eliminate the requirement of *a priori* specifying the dimensionality of the physical model describing a molecular complex’s kinetics. Here, we apply these methods to experimentally obtained datasets with samples illuminated by time-invariant laser intensities. In particular, we study interactions of IDPs.

## 1 Terminology Convention

To be consistent throughout our three part manuscript, we precisely define some terms as follows

1. a macromolecular complex under study is always referred to as a *system*,
2. the configurations through which a system transitions are termed *system states*, typically labeled using *σ*,
3. FRET dyes undergo quantum mechanical transitions between *photophysical states*, typically labeled using *ψ*,
4. a system-FRET combination is always referred to as a *composite*,
5. a composite undergoes transitions among its *superstates*, typically labeled using *ϕ*,
6. all transition rates are typically labeled using λ,
7. the symbol *N* is generally used to represent the total number of discretized time windows, typically labeled with *n*, and
8. the symbol *w_n_* is generally used to represent the observations in the *n*-th time window.

## 2 Introduction

Single molecule Förster Resonance Energy Transfer (smFRET) experiments are widely used [1] to study molecular kinetics across timescales on both stationary [2–5] and freely diffusing molecules [6]. These timescales include faster events, below the micro-to millisecond timescales, including domain rotations, configurational kinetics of disordered proteins, protein folding, protein-protein interactions, all the way to slower events, such as misfolding and refolding events, occurring on minute and even hour long timescales [7].

In a typical experiment we consider herein, a continuous wave (CW) laser illuminates a sample with a beam of constant intensity and power over a period of time. CW sources are common as they are both cheaper and technically simpler to implement in an experimental setup than their pulsed counterparts [8, 9] that we explore in our third companion manuscript [10]. However, as compared to pulsed sources, a disadvantage lies in the increased photon flux through the sample which can accelerate photodamage [11].

While pulsed illumination can significantly reduce sample photobleaching and phototoxicity [12] and more readily reveals excited state lifetimes of fluorophores, in practice it is restricted to analyzing one (time-stamped) photon per interpulse period. This in turn limits the data acquisition rate and sets a bound on the temporal resolution of the kinetics we may deduce from pulsed single photon arrival.

By contrast, continuous illumination avoids this problem, by allowing a larger number of photons to be detected in the time that would normally be considered an interpulse period in pulsed illumination [13]. The cost then comes at the loss of direct knowledge of excited state lifetime which can, with difficulty and high uncertainty, then be decoded from photon-antibunching statistics if required [14] as shown in the first companion manuscript [15].

It is common practice to analyze photon arrival data to extract kinetics under continuous illumination by binning the data and subsequently using hidden Markov models (HMMs) [16–19]. As noise distributions are better characterized in unprocessed data, it remains conceptually preferred, though more computationally costly, to use photon-by-photon methods [13, 14, 20–24]. Indeed, photon-by-photon methods can be used to learn both photophysical and system transition rates directly from the detected photon colors and interphoton arrival times. Additionally, this has the benefit of avoiding averaging kinetics that may occur when binning data [17].

Currently available methods to analyze smFRET data in a photon-by-photon manner [13, 20] rely on the foundational works of Gopich and Szabo [13, 14, 25], where the likelihood is taken as the product of as many generator matrix exponentials as there are photons in a FRET trace. Such a generator matrix constitutes transition rates encoding the kinetics of the system-FRET composite [15].

When analyzing smFRET data, of particular interest is the dimensionality of this generator matrix determined by the number of system states. In all existing analyses, the dimensionality is fixed by hand *a priori* and the transition rates are then learned as point estimates using maximum likelihood methods.

Yet point estimates can be biased. In fact, limited data, lack of temporal resolution to estimate very fast kinetics [15], and noise, all contribute to bias [26] in addition to a flattening of possibly multimodal likelihoods [27, 28]. This motivates why we wish to operate in a Bayesian setting to learn distributions over the number of system states and transition rates, while incorporating unavoidable noise sources such as detector electronics and background.

For this reason, we developed a complete Bayesian nonparametric (BNP) framework in the first companion manuscript [15]. This framework incorporates many key complexities of a typical smFRET experimental setup, including background emissions, fluorophore photophysics (blinking, photobleaching, and direct acceptor excitation), instrument response function (IRF), detector dead time, and crosstalk.

Here, we delve deeper into this framework for the case of continuous illumination by exploring its utility in cases where the number of system states is unknown.

We first test the robustness of our nonparametric method and its software implementation BNP-FRET by analyzing synthetically generated data for kinetics varying from very slow to timescales as fast as the interphoton arrival times. We then apply our method to experimental smFRET data capturing interactions between intrinsically disordered protein (IDP) fragments [29, 30] relevant to signaling and regulation.

IDPs are of particular interest to nonparametric analyses as IDP’s lack of order and stability results in broader spectra of dominant FRET pair distances sensitive to their chemical environment. In particular, we study interactions between the nuclear-coactivator binding domain (NCBD) of a CBP/p300, *i.e*., transcription coactivator and the activation domain of SRC −3 (ACTR) under varying chemical conditions affecting their coupled folding and binding reaction rates [29–31]. We use a single FRET pair under continuous illumination to observe the possible physical configurations (system states) of the NCBD-ACTR complex. Further, we report new bound/transient system states for the NCBD P20A mutation, not observed using previous point estimation techniques [30].

## 3 Forward Model and Inference Strategy

For the sake of completeness, we begin with relevant aspects of the methods presented in the first companion manuscript [15], including the likelihood needed in Bayesian inference, and our parametric and nonparametric Markov Chain Monte Carlo (MCMC) samplers.

An smFRET experiment involves at least two single photon detectors collecting information on stochastic arrival times. We denote these arrival times with

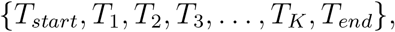

in detection channels

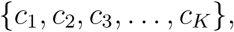

for a total number of *K* photons. In this representation above, *T_start_* and *T_end_* are the experiment’s start and end times, respectively.

Using this dataset, we would like to infer parameters governing a system’s kinetics. That is, the number of system states *M_σ_* and the associated transition rates *λ_σ_i_ →σ_j__*, as well as *M_ψ_* photophysical transition rates 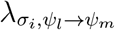 corresponding to each system state *σ_i_*. Here, *σ_i_* ∈ {*σ*_1_, …, *σ_M_σ__* } and *ψ_l_* ∈ {*ψ*_1_, …, *ψ_M_ψ__*} are the system states and photophysical states, respectively. These rates populate a generator matrix **G** of dimension *M_ϕ_* = *M_σ_* × *M_ψ_* now representing transitions among composite superstates, *ϕ_i_* ≡ (*σ_j_*, *ψ_k_*) where *i* = (*j* – 1)*M_ψ_* + *k* (see the first companion manuscript for details [15] on the structure of such a matrix). This matrix governs the evolution of the system-FRET composite via the master equation

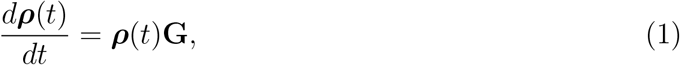

as described in Sec. 2.3 of the first companion manuscript [15]. Here, ***ρ***(*t*) is a row vector populated by probabilities for finding the composite in a given superstate at time *t*.

In estimating these parameters, we must account for all sources of uncertainty present in the experiment, such as shot noise and detector electronics. Therefore, we naturally work within the Bayesian paradigm where the parameters are learned by sampling from probability distributions over these parameters termed posteriors. Such posteriors are proportional to the product of the likelihood, which is the probability of the collected data ***w*** given the physical model, and prior distributions over the parameters as follows

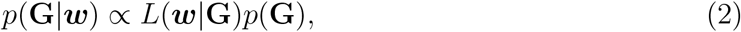

where ***w*** constitutes the set of all observations, including photon arrival times and detection channels.

To construct the posterior, we begin with the likelihood

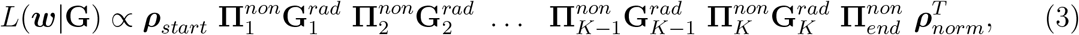

derived in Sec. 2.3 of the first companion manuscript. Here, 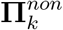 and 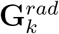 are the non-radiative and radiative propagators, respectively. Furthermore, ***ρ**_start_* is computed by solving the master equation assuming the system was at steady-state immediately preceding the time at which the experiment began. That is, we solve

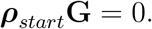

Next, assuming that the transition rates are independent of each other, we can write the associated prior as

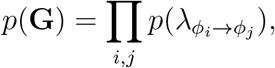

where we choose Gamma prior distributions over individual rates. That is,

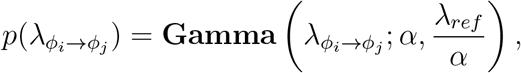

to guarantee positive values. Here, *ϕ_i_* represents one of the *M_ϕ_* superstates of the system-FRET composite collecting both the system and photophysical states as described in Sec. 2.2. Furthermore, *α* and λ*_ref_* are parameters of the Gamma prior.

In what follows, we first assume that the number of system states are known and will describe an inverse strategy that uses the posterior above to learn only transition rates. Next, we generalize our model to a nonparametric case accommodating more practical situations with unknown system state numbers. We do so by assuming an infinite dimensional system state space and making the existence of each system state itself a random variable.

### 3.1 Inference Procedure: Parametric Sampler

Now, with the posterior defined, we prescribe a sampling scheme to learn distributions over all parameters of interest, namely, transitions rates populating **G** and the number of system states. However, our posterior in Eq. 2 does not assume a form amenable to analytical calculations. Therefore, we employ Markov Chain Monte Carlo (MCMC) techniques to draw numerical samples.

Particularly convenient here is the Gibbs algorithm that sequentially and separately generates samples for individual transition rates in each MCMC iteration. This requires us to first write the posterior in Eq. 2 using the chain rule as follows

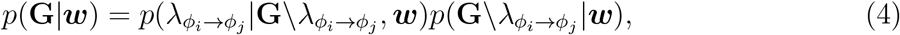

where the backslash after **G** indicates exclusion of the subsequent rate parameter. Furthermore, the first term on the right hand side is the conditional posterior for the individual rate λ*_ϕ_i_→ϕ_j__*. The second term in the product is a constant in the corresponding Gibbs step as it is independent of λ*_ϕ_i_→ϕ_j__*. Similarly, the priors *p*(**G**\λ*_ϕ_i_→ϕ_j__*) for the rest of the rate parameters on the right hand side of Eq. 2 are also considered constant. Equating the right hand sides of Eqs. 2 & 4 then allows us to write the following conditional posterior for λ*_ϕ_i_→ϕ_j__* as

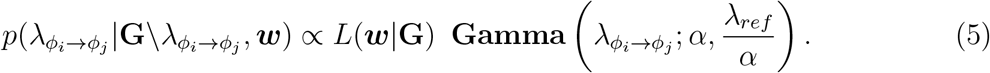

Since the conditional posterior above does not take a closed form that allows for direct sampling, we use the Metropolis-Hastings (MH) step [32–34] where new samples are drawn from a proposal distribution *q* and accepted with probability

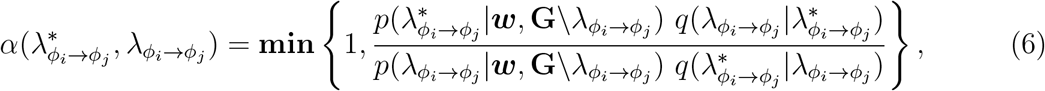

where the asterisk denotes proposed rate values from the proposal distribution *q*.

Now, to generate an MCMC chain of samples, we first initialize the chains for all transition rates λ*_ϕ_i_→ϕ_j__*, by randomly drawing values from their corresponding prior distributions. We then successively iterate across each transition rate in each new MCMC step and draw new samples from the corresponding conditional posterior using the MH criterion.

In the MH step, a convenient choice for the proposal is a Normal distribution leading to a simpler formula for the acceptance probability in Eq. 6. This is due to its symmetry resulting in 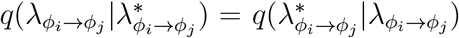. However, a Normal proposal distribution would allow forbidden negative transition rates, leading to automatic rejection in the MH step and thus inefficient sampling. Therefore, it is more convenient to propose new samples using a Normal distribution in logarithmic space to allow exploration along the full real line as follows

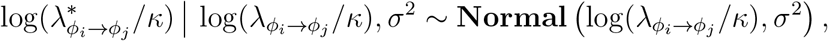

where *κ* = 1 is an auxiliary parameter in the same units as λ*_ϕ_i_→ϕ_j__* introduced to obtain a dimensionless quantity within the logarithm.

The transformation above requires introduction of Jacobian factors in the acceptance probability as follows

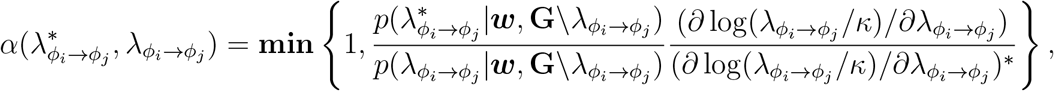

where the derivatives represent the Jacobian and the proposal distributions are canceled by virtue of using a Normal distribution.

The acceptance probability above depends on the difference of the current and proposed values for a given transition rate. This difference is determined by the covariance of the Normal proposal distribution *σ*^2^ which needs to be tuned for each rate individually to achieve an optimum performance of the BNP-FRET sampler, or equivalently approximately one-third acceptance rate for the proposals [35].

In our case, where the smFRET traces analyzed contain about 10^5^ photons, we found it prudent to make the sampler alternate between two sets of variances at every MCMC iteration, 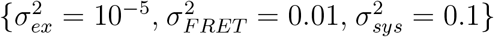 and 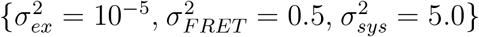, for the excitation rates, FRET rates, and system transition rates. This ensure that the sampler is quickly able to explore values at different orders of magnitude.

Intuitively, these covariance values in the proposal distributions above would ideally scale with the relative widths of the conditional posteriors for these parameters (in log-space) if the approximate width could be estimated. Since posterior widths depend on the amount of data used, an increase in the number of photons available in the analysis would require a correspondingly smaller variance.

### 3.2 Inference Procedure: Nonparametric BNP-FRET Sampler

Here, we first, briefly summarize our inference procedure described in Sec. 3.1 & 3.2.1 of the first companion manuscript [15] for ease of reference.

In realistic situations, the system state space’s dimensionality is usually unknown as molecules under study may exhibit complex and unexpected behaviors across conditions and timescales. Consequently, the dimensionality *M_ϕ_* of the generator matrix **G** is also unknown, and must be determined by adopting a BNP framework.

In such a framework, we assume an infinite set of system states and place a binary weight, termed load, on each system state such that if it is warranted by the data, the value of the load is realized to one. Put differently, we must place a Bernoulli prior on each candidate state (of which there are formally an infinite number) [36, 37]. In practice, we learn distributions over Bernoulli random variables *b_i_* that activate/deactivate different portions of the full generator matrix as (see Sec. 3.2.1 of the first companion manuscript [15])

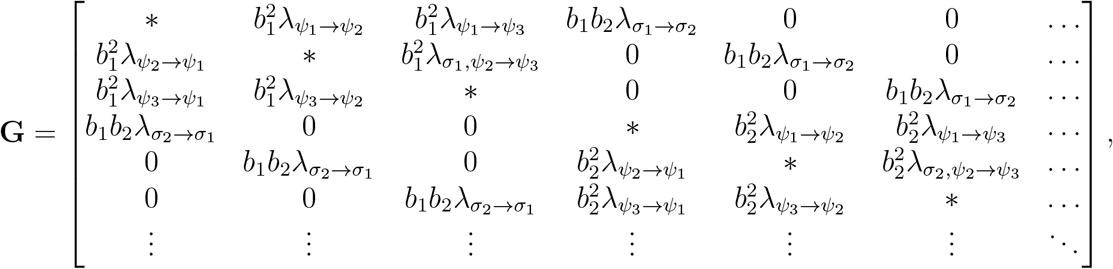

where active loads are set to 1, while inactive loads are set to 0. Furthermore, * represents negative row-sums. Finally, the number of active loads provides an estimate of the number of system states warranted by a given dataset.

As we have introduced new variables we wish to learn, we upgrade the posterior of Eq. 2 to incorporate the full set of loads, **b** = {*b*_1_, *b*_2_, …, *b*_∞_}, as follows

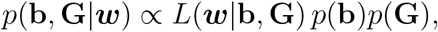

where we assume that all parameters of interest are independent of each other.

As in the parametric sampler presented in the previous subsection, we generate samples from the nonparametric posterior above using Gibbs sampling. That is, we first initialize the MCMC chains for loads and rates by drawing random samples from their priors. Next, to construct the chains, we iteratively draw samples from the posterior in two steps: 1) sequentially sample all rates using the MH procedure; then 2) loads by direct sampling, from their corresponding conditional posteriors (as described in Sec. 3.2.1 of the first companion manuscript [15]). Since step (1) is similar to the parametric case, we only focus on the second step in what follows.

To generates samples for load *b_i_*, the corresponding conditional posterior is given by [38]

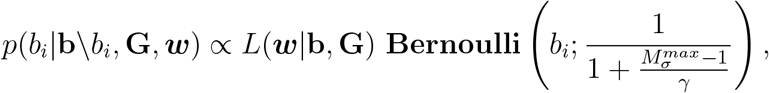

where the backslash after **b** indicates exclusion of the following load. We may set the hyperparameters 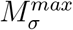, the maximum allowed number of system states used in computations, and *γ*, the expected number of system states based on simple visual inspection of the smFRET traces.

Now, the conditional posterior in the equation above is discrete and describes the probability for the load to be either active or inactive, that is, it is itself a Bernoulli distribution as follows

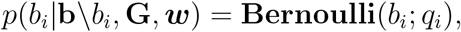

where

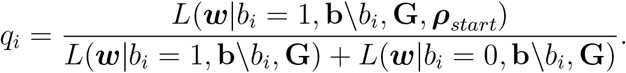

The simple form of this posterior is amenable to direct sampling. In the end, the chain of generated samples can be used for subsequent statistical analysis.

## 4 Results

In this section, we first demonstrate the robustness of our BNP-FRET sampler by investigating the effects of excitation rate on the distributions over transitions rates and system state numbers. Once we have illustrated the BNP-FRET sampler’s performance on synthetic data, we apply it to estimate the number of system states along with associated escape rates from publicly available experimental data for a complex involving instrinsically disorder proteins (ACTR-NCBD). We compare our results with reported literature values [29, 30].

### 4.1 Resolution of Timescales Given Excitation Rate: Nonparametrics

To demonstrate the performance of our BNP-FRET sampler over a range of timescales given a fixed excitation rate, we follow the same approach as presented in the first companion manuscript (see Sec. 4.1) [15]. That is, we generate four synthetic smFRET traces containing *K* = 2 million photons each for a biomolecular complex with three system states, {*σ*_1_, *σ*_2_, *σ*_3_}. The kinetic scheme for this system is a generalization of the example presented in the first companion manuscript [15] (brown boxes) with two system states.

Now, to synthesize smFRET traces, we fix the excitation rate to λ*_ex_* = 10 ms^−1^ and FRET efficiencies *ε_FRET_* to 0.09, 0.5, and 0.9 for the three system states, respectively, motivated by experiments in [30]. The remaining parameters are the system transition rates λ_*σ_i_*→*σ_j_*_, varied across datasets to test our BNP-FRET sampler over a wide range of timescales ranging from a thousand times longer than the average interphoton arrival time (1/λ_*ex*_) to as short as the average interphoton arrival time itself (representing an extreme case). We do not probe kinetics any faster because the excitation rate does not provide enough temporal resolution for resolving system transitions in this regime, as demonstrated in the first manuscript (see Sec. 4.1).

We start the analysis by applying our BNP-FRET sampler to learn the number of system states for the case with slowest escape rates, *i.e*., the sum of all transition rates out of a given system state. These escape rates are λ*_esc_* = 0.01, 0.02, and 0.03ms^−1^. We show that our BNP-FRET sampler can correctly learn the number of system states and the associated escape rates and FRET efficiencies; see Fig. 1(a) and Fig. 2(a).

**Figure 1:**
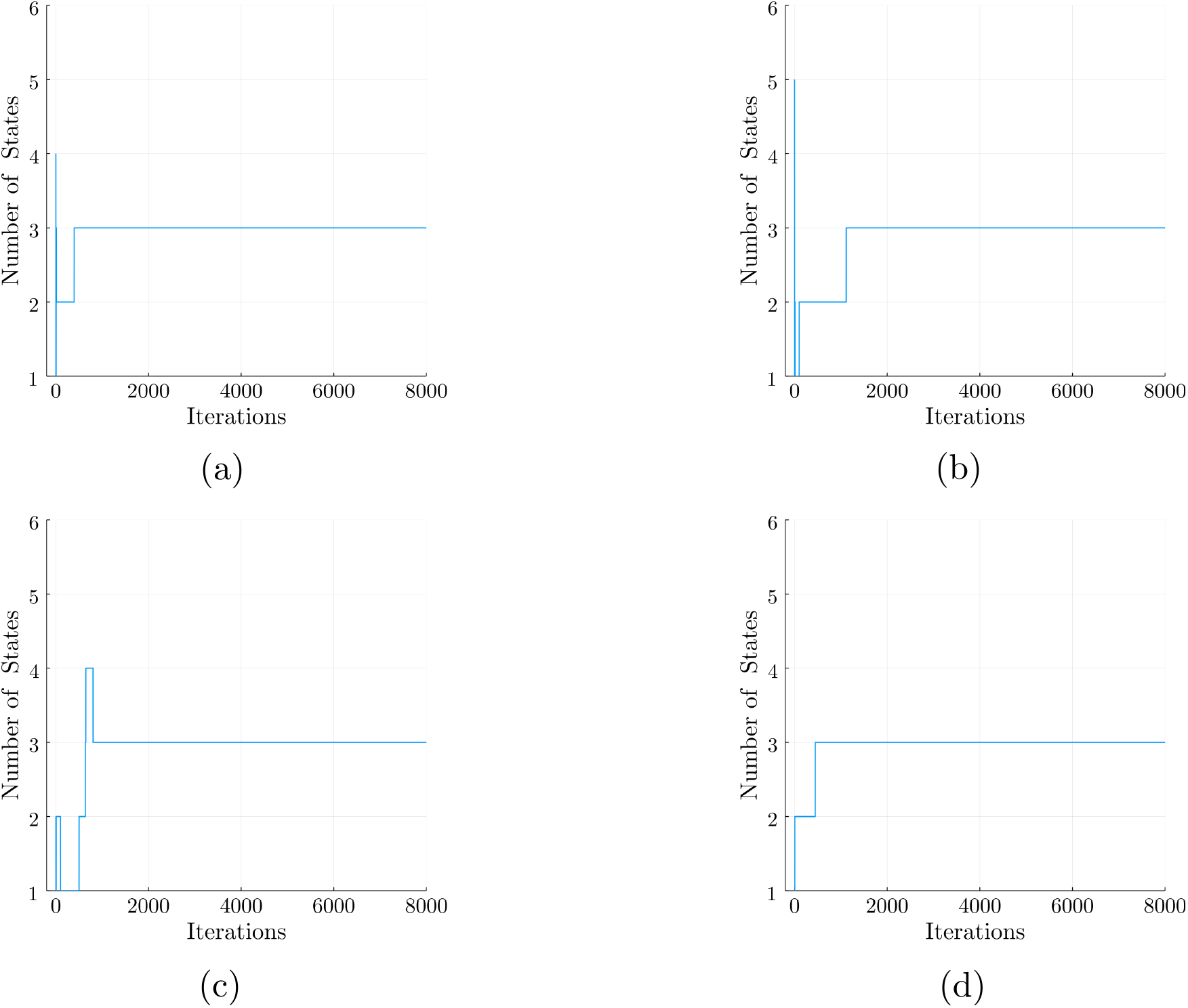
MCMC chains generated by the BNP-FRET sampler for the number of system states. The synthetic smFRET datasets used to generate these chains assume uniform excitation rate of 10 ms^−1^ and FRET efficiencies of 0.09, 0.5, and 0.9, for a three state system. However, the system’s escape rates for all three states become faster by a factor of 10 as we move from panel (a) to (d). That is, in the slowest case, we use escape rates of 0.01, 0.02, and 0.03 ms^−1^ for the three system states, while in the fastest case kinetics are as fast as the excitation rate itself. Our method converges to the correct number of system states for each dataset. As we will see later, the rates become more difficult to estimate for panel (d) which we consider to be the point at which the method breaks down.

**Figure 2:**
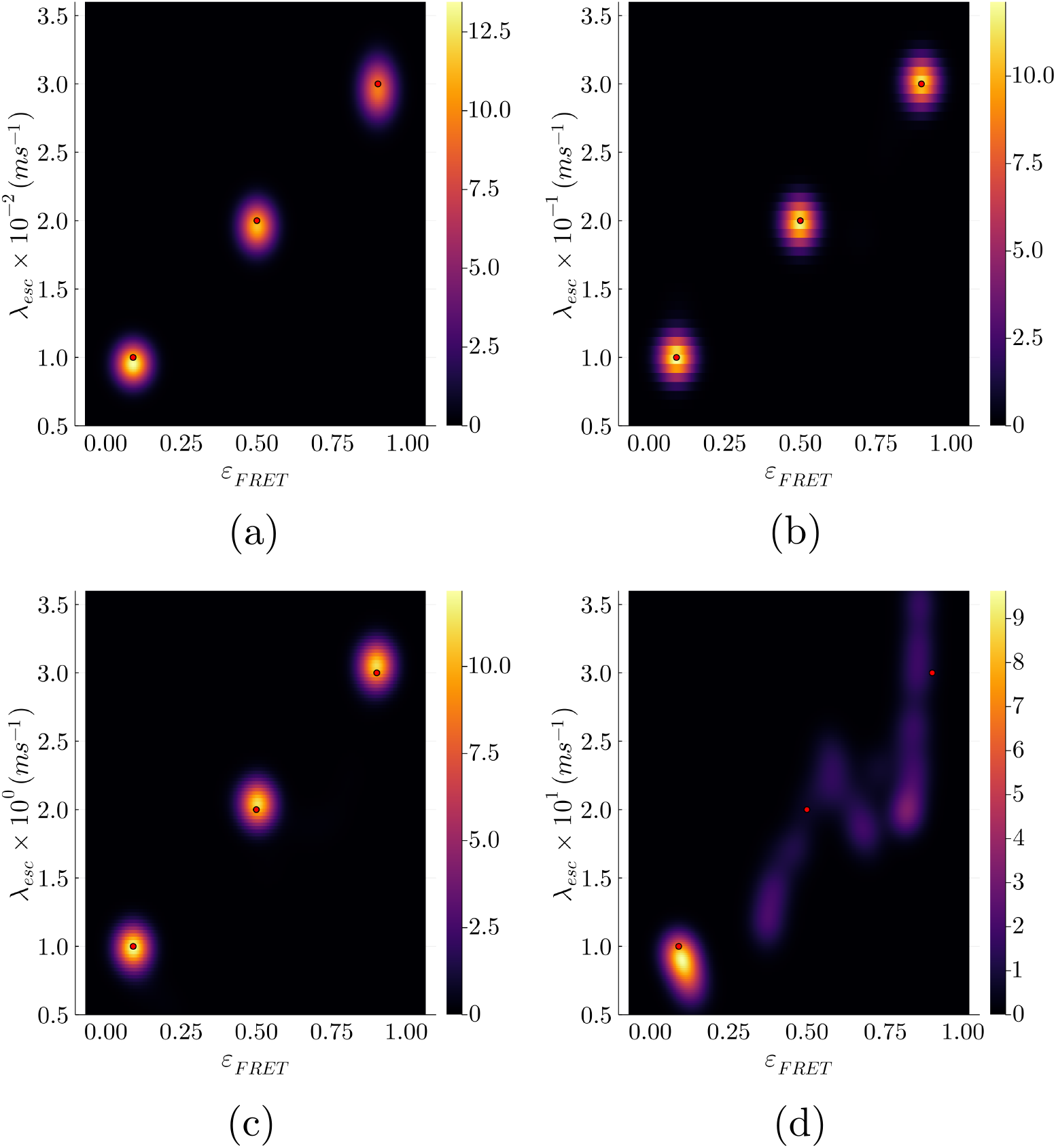
Learned bivariate posterior for the escape rates λ_*esc*_ and FRET efficiencies *ε_FRET_* from synthetic data also used in Fig. 1. Going from panels (a) to (d), we speed up the kinetics (escape rates) by a factor of 10 each time leading to a gradual loss of temporal resolution needed to identify system transitions. The ground truth is shown with the red dots. The estimates for escape rates and FRET rates in panels (a) to (c), have less than 10% errors. However, as seen in panel (d), the excitation rate does not provide enough temporal resolution to resolve system transitions occurring at interphoton arrival time-scales, resulting in large errors in the parameter estimates. The estimated escape rates in panel (d) are 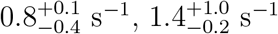, and, 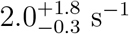 with very large uncertainties (95% confidence intervals). We have smoothed the posterior distributions here using KDE for visualization purposes only.

Next we analyze, one-by-one, datasets generated using escape rates that are 10 times faster in each subsequent dataset. BNP-FRET deduces the correct number of system states in all cases (see Fig. 2a-c), however the determination of the rates begins to fail in panel (d) of Fig. 2.

The failure to estimate escape rates approximating the excitation rate can also be predicted using a “photon budget index” defined in the first companion manuscript [15] Sec. 4.1 as

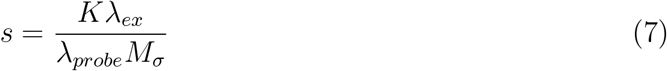

where *K* and λ_*probe*_ are, respectively, the photon counts and the escape rate to be probed. Plugging the parameter values associated to the dataset shown in both Fig. 1(d) and 2(d) with three escape rates, *i.e., K* = 2 × 10^6^, *M_σ_* = 3, λ_*ex*_ = 10ms^−1^ and λ_*probe*_ = λ_*esc*_ = 10 – 30ms^−1^, into the above equation, we obtain *s* = 2/3 × 10^6^, 2/6 × 10^6^ and 2/9 × 10^6^. The index obtained for λ_*probe*_ = 10ms^−1^ is on par with the threshold of *s*_thresh_ = 10^6^ derived in the first companion manuscript in [15] Sec. 4.1 where the sampler had available sufficient information to drawn an accurate inference. By contrast, moving to the larger escape rates of λ_*esc*_ = 20, 30ms^−1^ the photon budget indices obtained are much smaller than the threshold and the sampler starts failing due to lack of information. To be more precise, our sampler is capable of learning any escape rates, even those larger than excitation rate, given sufficient photons. As this is counter intuitive, we note that the excitation rate is an average value and there are often photons detected with interphoton intervals much smaller than 1/λ_*ex*_. As such, given long photon traces, there are always enough photons with small interphoton intervals to learn faster escape rates (and indeed to learn excited state lifetimes as we show in the first companion manuscript [15]) that would otherwise evade binned photon analysis methods [39].

### 4.2 Analysis of Experimental Data: NCBD-ACTR Interactions

Here, we apply our BNP-FRET sampler to two datasets probing the interactions between partner IDPs, NCBD and ACTR, under different conditions [29, 30]. Precise knowledge of binding and unbinding reactions of such proteins is of fundamental importance toward understanding how they regulate expression of their target genes.

Methods that have been used in the past [29, 30] to analyze smFRET traces from experiments on NCBD-ACTR interaction assumed a fixed number of system states to obtain maximum likelihood point estimates for transition rates. In addition, these methods bin photons to mitigate computational expense. However, given the inherently unstructured and flexible nature of IDPs, fixing the dimensionality of the model *a priori* can be limiting and, as we will see, may bias analysis. Therefore, our nonparametric method which places no constraints on the number of system states while incorporating all major noise sources, is naturally suited.

In the following subsections, we first analyze data for a system where an immobilized ACTR labeled with a Cy3B donor interacts with an NCBD labeled with a CF680R acceptor in the presence of ethylene glycol (EG), 36% by volume, in order to more closely mimic cellular viscosity [29]. Here, the binding of NCBD to ACTR is monitored in smFRET experiments using a confocal microscope setup. Next, we analyze data for a system in a buffer without EG, and therefore with faster kinetics. Here an immobilized ACTR interacts with a freely-diffusing mutated NCBD (P20A) [30].

To acquire both experimental FRET datasets containing about 200000 photons each, laser powers of 0.5 *μ*W and 0.3 *μ*W were used leading to excitation rates varying from 3000 to 11000 s^−1^ in the confocal region depending on where the immobilized sample lies with respect to the center of the excitation laser beam.

Moreover, we are provided a calibrated route correction matrix (RCM) by the authors of [29, 30] to account for spectral crosstalk, and relative detection efficiencies of donor and acceptor channels. We defined such an RCM in Sec. 2.4.1 of the first companion manuscript [15] and specify it for each dataset separately in the following subsections.

Finally, by contrast to the first companion manuscript [15] we ignore the IRF. The latter typically acts over a period of hundreds of picoseconds. As such, it is immaterial on the seconds timescale over which system transitions occur. Moreover, the background values vary for each dataset, and are therefore precalibrated, independently, for each dataset in the corresponding sections.

Now, with all experimental details at hand, we proceed to analyze the experimental data using our BNP-FRET sampler.

#### 4.2.1 Immobilized ACTR in 36% EG

Binding of NCBD to ACTR leads to the formation of a stable and ordered complex in the presence of EG. In addition, when two fluorescent dyes labeling the IDPs come in close proximity, we expect FRET interactions. Therefore, bound and unbound system states of the NCBD-ACTR complex correspond to high and low FRET efficiency signals, respectively.

For the analysis of the collected smFRET data from such a complex, we must take into account all sources of noise such as crosstalk and background. The crosstalk/detection efficiency values are computed from the RCM given by the authors of [29] as

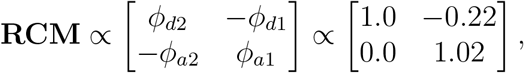

where channels 1 & 2 are, respectively, designed to receive acceptor and donor photons. Furthermore, *ϕ_ai_* and *ϕ_di_*, respectively, denote probabilities of acceptor and donor photons being registered by channel *i*. Adopting the same normalization convention for the RCM as in the first companion manuscript [15] (see Example V) gives the following values for the effective crosstalk factors as

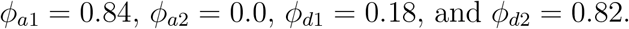

As such, these values imply that approximately 18% of the emitted donor photons are detected in the acceptor channel due to crosstalk. Furthermore, only 84% of emitted acceptor photons are detected in the acceptor channel, and acceptor photons do not suffer any crosstalk.

We must also incorporate precalibrated background rates for donor and acceptor channels given as 0.283 s^−1^ and 0.467 s^−1^, respectively [29].

With all such corrections applied, our BNP-FRET sampler now predicts two system states; see Fig. 3. The system state with the lowest FRET efficiency of 0.0 corresponds to the unbound NCBD. The remaining system state with higher FRET efficiency of ≈ 0.7 coincides with the bound NCBD-ACTR complex configuration. The associated escape rates we obtain from our method for both of the system states are approximately 2.9 s^−1^ and 4.1 s^−1^ as seen in Fig. 3b. These results are consistent with results reported in supplementary table S1 of [29] with an average relative difference of ≈ 15%.

**Figure 3:**
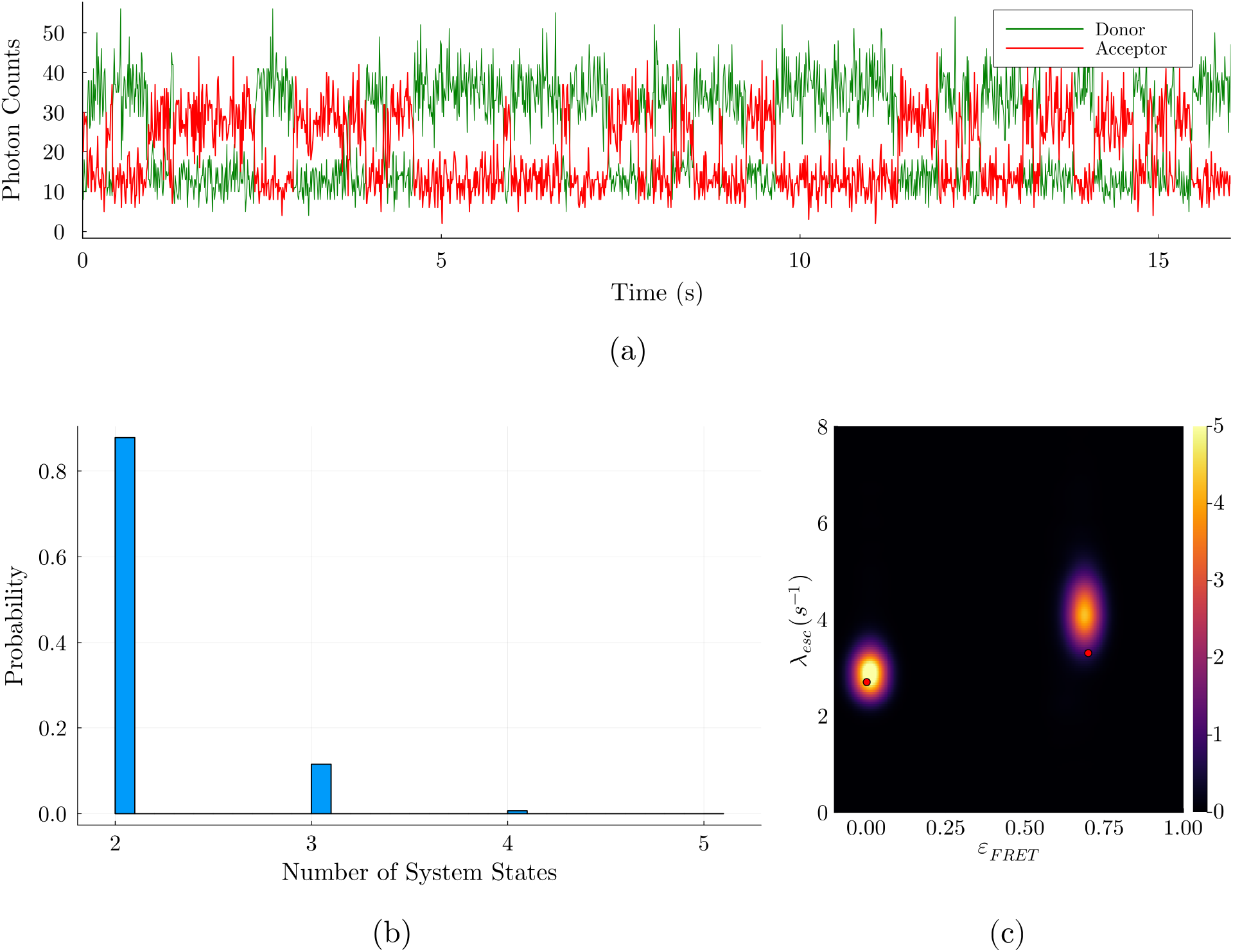
Results for NCBD-ACTR interactions in the presence of ethylene glycol (EG). Panel (a) shows the raw photon counts (bin width of 0.01s) recorded by the two detection channels during the experiment. In panel (b), we show a probability distribution for the number of system states estimated by the BNP-FRET sampler. The sampler spends a majority of its time in two system states with only a small relative probability ascribed to more states. In the posterior distribution for the escape rates and FRET efficiencies in panel (c), two distinct FRET efficiencies are evident with values of about 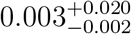 (unbound) and 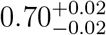 (bound), and corresponding escape rates of about 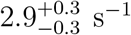 and 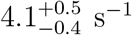. The red dots show results reported by [29] using maximum likelihood method. We have smoothed the distribution for demonstrative purposes only.

#### 4.2.2 Immobilized ACTR in buffer

Here, in the absence of EG, the viscosity of the solution is lowered [29], leading to faster system transitions representing a unique analysis challenge.

As in the previous subsection, from the RCM provided by the authors of [30] for the current dataset, we found crosstalk factors of *ϕ*_*a*1_ = 0.72, *ϕ*_*a*2_ = 0.0, *ϕ*_*d*1_ = 0.10, and *ϕ*_*d*2_ = 0.90. After correcting for these crosstalk/detection efficiency values and background rates of 0.312 s^−1^ and 1.561 s^−1^ for the donor and acceptor channels, respectively, our BNP-FRET sampler now predicts five system states (see Fig. 4(a)&(b)) with FRET efficiencies of 0.0, 0.72, 0.03, 0.28, and 0.92 approximately. Here, the first two system states with vanishingly small estimated FRET efficiencies, namely 0.0 and 0.03, most likely represent the same configuration where NCBD is diffusing freely away from the immobilized ACTR, leading to no FRET interactions. Various sources of noise in the dataset may have resulted in this splitting of the unbound system state. Furthermore, the system state with the FRET efficiency and escape rate of approximately 0.72 and 25.0 s^−1^, respectively, coincides with the previously predicted bound configuration found using a maximum likelihood method with a fixed number of system states [30]. We have compiled the learned transition rates (median values) in the generator matrix below (in s^−1^ units)

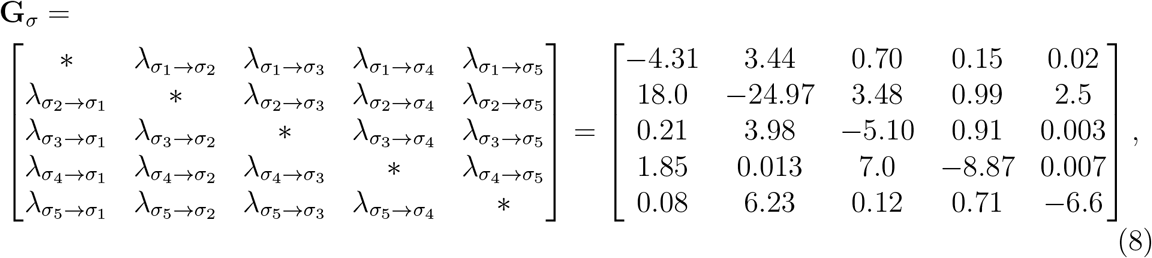

where the diagonal elements correspond to negative of the escape rate values. Furthermore, the steady-state populations/probabilities for these system states can be computed by solving ***ρ_steady_*****G**_*σ*_ = 0, resulting in

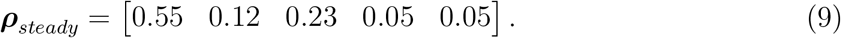

**Figure 4:**
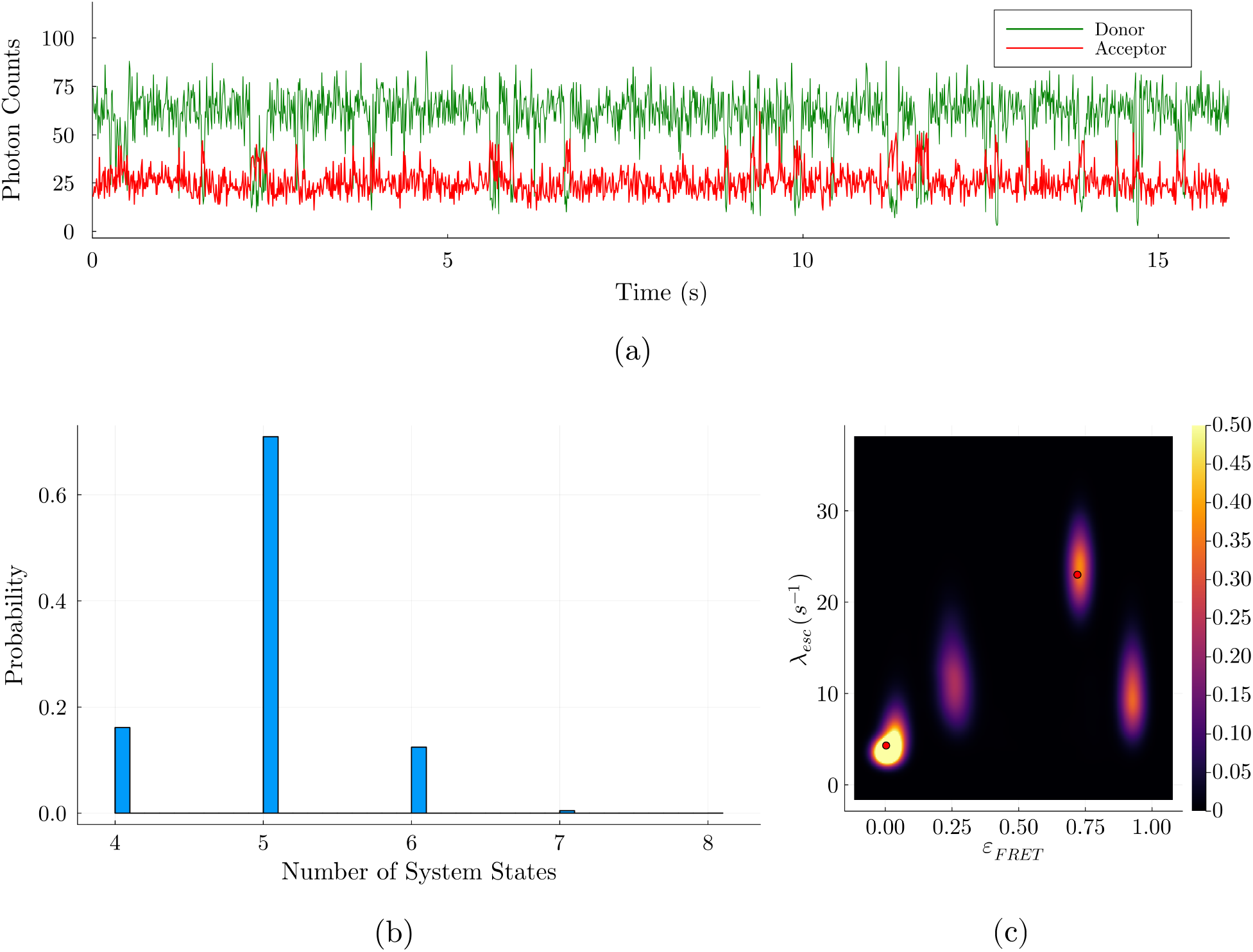
Results for NCBD-ACTR interactions in buffer, without EG. Panel (a) shows the raw photon counts (bin width of 0.01s) recorded by the two detection channels during the experiment. In panel (b), we show a probability distribution produced by the BNP-FRET sampler for the number of system states. Models with less than four system states in the histogram are not shown as we ascribe to them zero probability. Indeed, the most probable model contains five system states. Next, in panel (c) depicting the posterior distribution for the escape rates and FRET efficiencies, five distinct FRET efficiencies are evident with values of 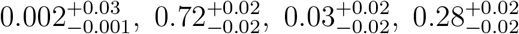, and 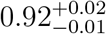 with corresponding escape rates of about 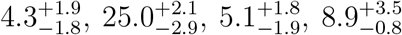, and 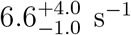. The first two system states with almost vanishing FRET efficiencies may represent the same unbound configuration with the small splitting likely arising from various sources of noise present in the dataset. The red dots show the results reported in [30] using maximum likelihood method.

Here, the two newly observed system states, with FRET efficiencies of 0.28 and 0.92 and corresponding escape rates of approximately 8.87 s^−1^ and 6.6 s^−1^, are bound configurations not previously detected [30] and deserve further attention. For instance, lower viscosity buffer (as compared to cases in the presence of EG) may allow the system to visit transient system states more readily under observation timescales [40, 41]. Additionally, steady-state probabilities for these new transient system states that we recover are indeed expectedly low (0.05 and 0.05) as compared to other system states of the NCBD-ACTR complex. Furthermore, IDPs interact in a complex manner with high possibility for residual secondary structures [42]. Competing parametric methods would need to posit a high number of system states *a priori* in order for their kinetics to be quantifiable. Finally, despite a difference in the estimate of the number of system states, our slower kinetics in the presence of EG are consistent with those of Ref. [29]. Direct comparison of escape rates across system states recovered by BNP-FRET versus Ref. [29] however is questionable on account of having recovered a different number of system states.

One way by which we may assure ourselves that these system states are not artefactually added by our computational algorithm (overfitting), is to analyze synthetic data generated under the same conditions (excitation rate, crosstalk, and background) as the experiment but with a ground truth of two system states. We can then ask whether the noise properties force our method to introduce artefactual states. Thus, we simulate a two system state model with the previously reported escape rates [30]) of 4.3 s^−1^ and 23.0 s^−1^ with corresponding FRET efficiencies of 0.0 and 0.8, and the same photon budget of 200000 photons. The results for the analysis of this synthetic dataset in Fig. 5(a)&(b) show no additional system states introduced by our method under this parameter regime suggesting the robustness of our findings for the experimental data.

**Figure 5:**
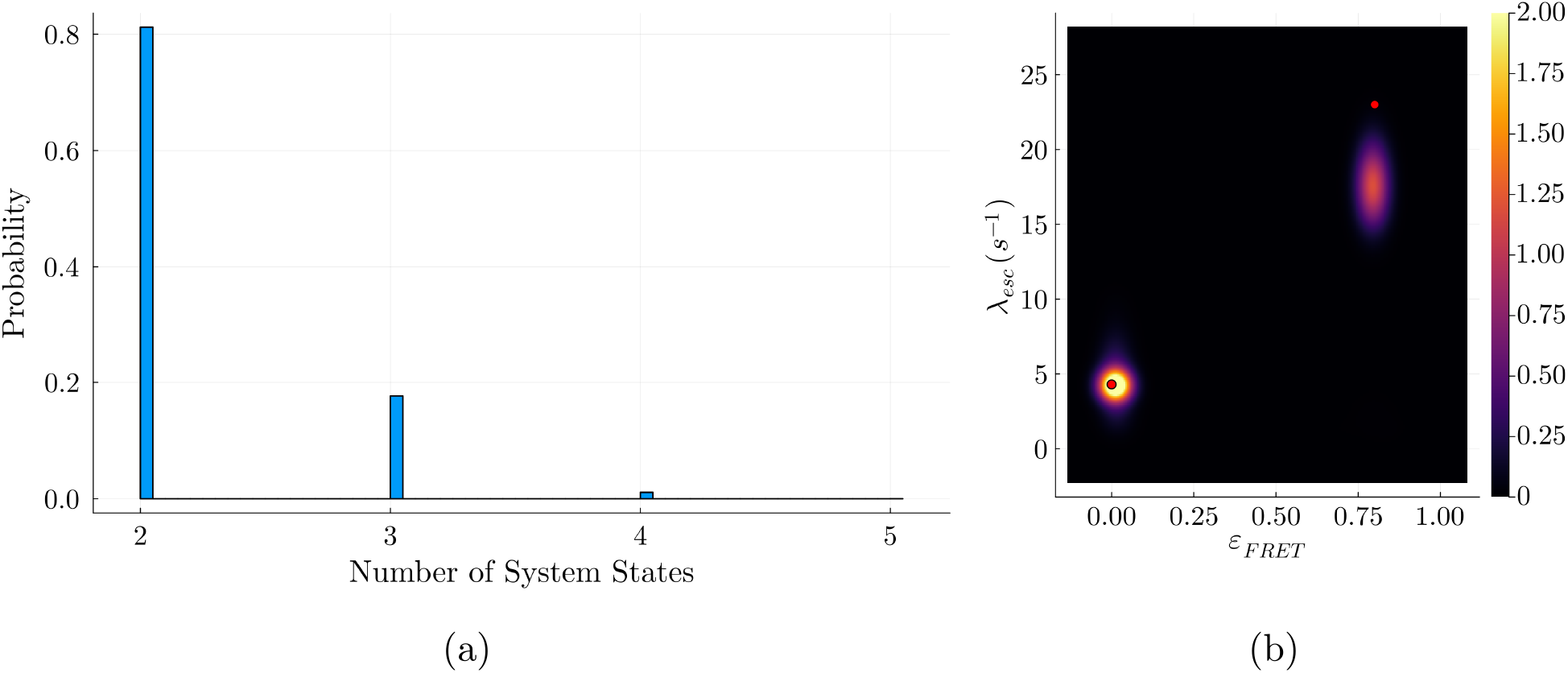
Robustness test using synthetic data with realistic noise parameters. The synthetic data here is generated under the same conditions (excitation rate, crosstalk, background, and photon budget) as the experiment whose results are shown in Fig. 4 with only two states as ground truth to see whether the multiple noise sources are likely to result in our method introducing spurious states (such as five states as seen in Fig. 4 using previously reported transitions rates [30]). In panel (a), we show the posterior produced by the BNP-FRET sampler for the number of system states. Fortunately, the most sampled model contains two system states, showing that noise sources do not introduce spurious states in this case. Small relative probability is ascribed to higher dimensional models. In the joint posterior distribution over the escape rates and FRET efficiencies in panel (b), two distinct FRET efficiencies are evident with values of 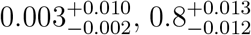 with corresponding escape rates of about 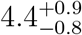 and 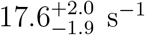. The red dots show the ground truth. The slight bias away from the ground truth results from high noise (background) in the data. The absence of additional system states suggests that the additional system states encountered in the experimental results are not artefactual.

Another way by which we may assure ourselves is by analyzing synthetically generated data for the four most distinct system states (on the basis of FRET efficiency) predicted by the BNP-FRET sampler for the experimental dataset. These system states correspond to FRET efficiencies of approximately 0.0, 0.72, 0.28, and 0.92 with associated escape rates of 4.31, 24.97, 8.87, and 6.6 s^−1^ as computed from the matrix in Eq. 8. We tested whether our sampler BNP-FRET underfits or overfits with regards to the estimated number of system states. As shown in Fig. 6, the most probable model predicted by the sampler has four system states, again verifying the robustness of our method.

**Figure 6:**
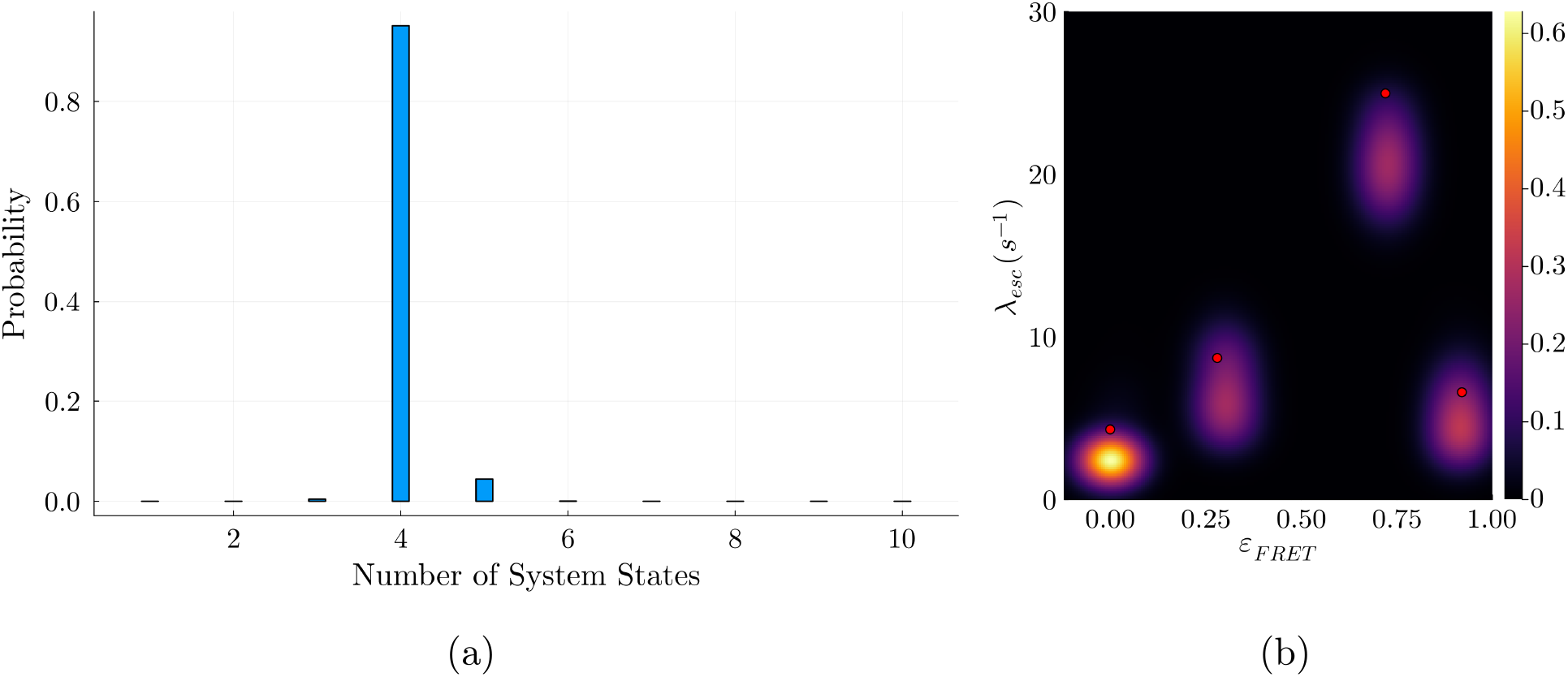
Second robustness test using synthetic data with realistic noise parameters. The synthetic data here is generated under the same conditions (excitation rate, crosstalk, background, and photon budget) as the experiment whose results are shown in Fig. 4 with four distinct system states (on the basis of FRET efficiency) as ground truth to see whether our sampler overfits or underfits with regards to the number of system states. These system states correspond to FRET efficiencies of 0.0, 0.72, 0.28, and 0.92 with associated escape rates of 4.31, 24.97, 8.87, and 6.6 s^−1^ as computed from the matrix in Eq. 8. In panel (a), we show the posterior produced by the BNP-FRET sampler for the number of system states. Fortunately, the most sampled model contains four system states, verifying the robustness of our method. Small probabilities are also ascribed to models with different numbers of system states. In the posterior distribution over the escape rates and FRET efficiencies in panel (b), four distinct FRET efficiencies are evident with values of 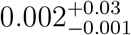, 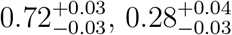 and 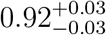 and corresponding escape rates of 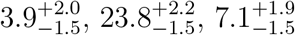, and 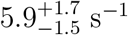. Here, the ground truth is shown with red dots. High noise from background results in the underestimates seen here.

## 5 Discussion

FRET techniques have been essential in investigating molecular interactions on nanometer scales, for instance, most recently in directly monitoring interaction of the SARS-COV2 virus spike protein with host receptors [43, 44]. Yet, the quantitative interpretation of smFRET data suffers from several issues including difficulties in estimating the number of system states, dealing with fast transition rates and providing uncertainties over estimates, particularly uncertainties over the number of system states [45, 46] originating from multiple noise sources.

Here, we implemented a general nonparametric smFRET data analysis framework presented in the first companion manuscript [15] to address the issues associated with smFRET data analysis acquired under continuous illumination. The framework developed can learn posterior distributions over the number of system states as well as the corresponding kinetics ranging from slow values all the way up to kinetic of events occurring on timescales approaching excitation rates. That is, our method propagates uncertainty over not only kinetic parameters but their associated models as well. This is especially significant in avoiding over-commitment to any one model when multiple models are almost equally probable given the data.

We benchmarked our method starting from synthetic data with three system states with a range of different timescales. We challenged our method by simulating data with kinetics as fast as the interphoton arrival times and correctly deduced the system state numbers even under such extreme conditions. We further assessed our method using experimental data acquired observing NCBD interacting with ACTR under different ethylene glycol (EG) concentrations that may impact the timescales at which the binding/unbinding reactions occur. In the previous point estimate methods [29, 30], two system states were assumed *a priori* for 0 and 36% EG concentrations. However, our nonparametric method predicts the number of system states and obtains two additional system states in the absence of EG (fast kinetics). This observation may be tied to the inherently unstable nature of the two IDPs under investigation [40].

A careful treatment of how experimental noise propagates into uncertainties over the number of system states and rates does come with associated computational cost. Other methods have managed to mitigate these costs by making approximations including: 1) assuming kinetics much slower than fluorophore excitation and relaxation rates [13, 14]; 2) assuming fast dye photophysics is completely irrelevant to the system transition rate and that FRET efficiency sufficiently identifies transitions between system states [14]; 3) ignoring detector effects and relegating other noise sources, such as background, to post-processing steps [13]; and, most popularly, 4) binning data [45, 47, 48]. For the general case without such approximations, however, the primary computation—the likelihood—remains expensive due to the required evaluation of many matrix exponentials. This cost can be mitigated in a number of ways by, for instance, computing likelihoods for several data traces in parallel. The scaling of the method is provided in the first companion manuscript [15].

The method described in this paper was developed for cases with discrete system state spaces. For continuous state spaces, both the likelihood and priors would require major modification in the spirit of Refs. [49, 50].

Our framework can accommodate different illumination modalities such as alternating laser excitation (ALEX) [51] to directly excited both donor and acceptor dyes by assuming nonzero direct excitation rates in the generator matrix. Indeed, direct excitation of the acceptor would further help in the simultaneous determination of crosstalk factors, detection efficiencies, and quantum yield of the dyes alongside kinetics.

## 6 Code availability

The BNP-FRET software package is available on Github at https://github.com/LabPresse/BNP-FRET

## 8 Acknowledgments

We thank Weiqing Xu and Dr. Zeliha Kilic for regular feedback and help, especially during the development of the nonparametrics samplers. We also thank Prof. Benjamin Schuler, Dr. Daniel Nettels, and Oliver Stach for regular feedback and providing the necessary experimental data. S.P. acknowledges support from the NIH NIGMS (R01GM130745) for supporting early efforts in nonparametrics and NIH NIGMS (R01GM134426) for supporting single-photon efforts. The majority of the computations were performed on the Agave and Sol supercomputers at ASU.

